# AliSim-HPC: parallel sequence simulator for phylogenetics

**DOI:** 10.1101/2023.01.15.524158

**Authors:** Nhan Ly-Trong, Giuseppe M.J. Barca, Bui Quang Minh

## Abstract

**Motivation:** Sequence simulation plays a vital role in phylogenetics with many applications, such as evaluating phylogenetic methods, testing hypotheses, and generating training data for machine-learning applications. We recently introduced a new simulator for multiple sequence alignments called AliSim, which outperformed existing tools. However, with the increasing demands of simulating large data sets, AliSim is still slow due to its sequential implementation; for example, to simulate millions of sequence alignments, AliSim took several days or weeks. Parallelization has been used for many phylogenetic inference methods but not yet for sequence simulation.

**Results:** This paper introduces AliSim-HPC, which, for the first time, employs high-performance computing for phylogenetic simulations. AliSim-HPC parallelizes the simulation process at both multi-core and multi-CPU levels using the OpenMP and MPI libraries, respectively. AliSim-HPC is highly efficient and scalable, which reduces the runtime to simulate 100 large alignments from one day to 9 minutes using 256 CPU cores from a cluster with 6 computing nodes, a 162-fold speedup.

**Availability and implementation:** AliSim-HPC is open source and available as part of the new IQ-TREE version v2.2.2.2 at https://github.com/iqtree/iqtree2/releases with a user manual at http://www.iqtree.org/doc/AliSim.

## Introduction

Phylogenetic inference is an important problem in bioinformatics, which aims to reconstruct a phylogenetic tree that describes the evolutionary relationship among a set of organisms (Felsenstein, 2004; Lemey *et al*., 2009). Typical phylogenetic inference methods require a multiple sequence alignment (MSA) containing DNA or amino-acid sequences as input and return a phylogenetic tree and a substitution model as output (Fig. 1A). In a phylogenetic tree, tips (leaves) represent the organisms in the MSA, internal nodes denote the extinct common ancestors. A substitution model is typically a Markov process that describes the rates of changes between nucleotides (for DNA sequences) or amino acids (for protein sequences).

**Fig. 1.**
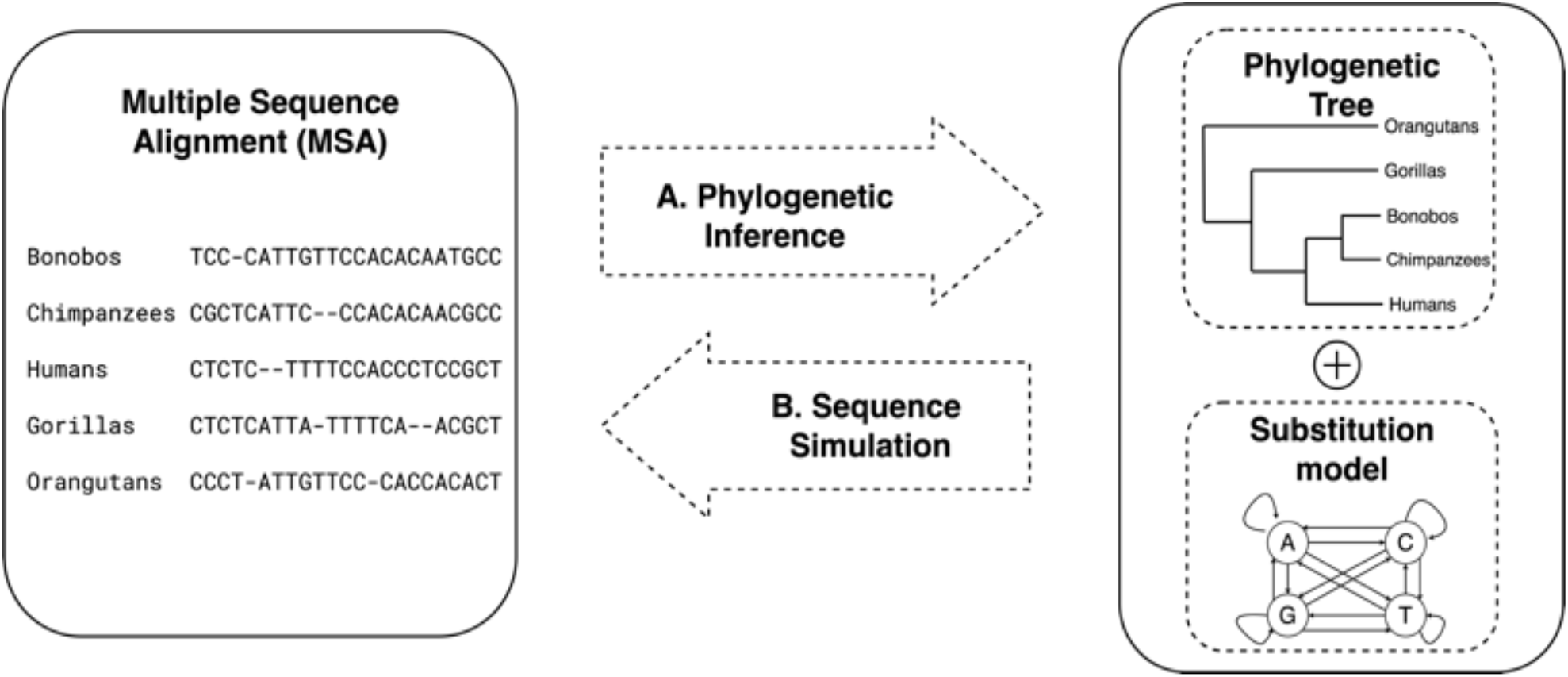
Phylogenetic inference (A) to infer a phylogenetic tree and a substitution model from an input multiple sequence alignment; and Sequence simulation (B) to generate multiple sequence alignment(s) from a phylogenetic tree and a substitution model.

Sequence simulation is an inverse problem of phylogenetic inference: we want to simulate MSAs from a phylogenetic tree and a substitution model (Fig. 1B). Simulated data has many applications in phylogenetics, such as evaluating phylogenetic methods (Garland *et al*., 1993; Kuhner and Felsenstein, 1994; Tateno *et al*., 1994; Huelsenbeck, 1995), testing hypothesis (Goldman, 1993; Adell and Dopazo, 1994; Schoeniger and von Haeseler, 1999), and more recently, generating data for training new machine learning methods (Abadi *et al*., 2020; Leuchtenberger *et al*., 2020; Ling *et al*., 2020; Suvorov *et al*., 2020).

We recently introduced a new sequence simulator, AliSim (Ly-Trong *et al*., 2022), which outperformed existing simulators (*i*.*e*., Seq-Gen (Rambaut and Grassly, 1997), Dawg (Cartwright, 2005), INDELible (Fletcher and Yang, 2009), and phastSim (De Maio et al., 2022)) regarding both running time and memory footprint. However, many applications require simulations of a vast number of large MSAs. For example, training new machine learning applications for phylogenetics (Suvorov and Schrider, 2022; Burgstaller-Muehlbacher *et al*., 2021; Abadi *et al*., 2020; Suvorov *et al*., 2020) requires millions of simulated MSAs. Due to its sequential implementation, AliSim becomes very slow, taking several days or weeks to simulate millions of alignments. Parallelization has been widely used for phylogenetic inference methods (Minh *et al*., 2020; Kozlov *et al*., 2015, 2019; Bouckaert *et al*., 2014; Morel *et al*., 2019; Altekar *et al*., 2004) but has not yet been employed for sequence simulation.

In this paper, we introduce AliSim-HPC, a high-performance computing version of AliSim. AliSim-HPC parallelizes the simulations at both multi-core and multi-CPU levels using OpenMP (Chapman *et al*., 2007) and the Message Passing Interface (MPI) (Gropp *et al*., 1998), respectively. We first present two multi-threading algorithms to parallelize the simulation of a single (large) alignment with the OpenMP library. Next, we utilize the MPI library to parallelize the simulation of many alignments across distributed CPUs. We can thus deploy AliSim-HPC that combines OpenMP and MPI on a high-performance computing cluster with many nodes. We note that the proposed algorithms are generally applicable to shared and distributed memory paradigms. We only chose OpenMP and MPI because these two libraries have already been used in IQ-TREE.

AliSim-HPC shows an excellent scaling behavior: it reduces the simulation time of 100 large alignments from one day to 9 minutes by using 256 CPU cores (162-fold speedup). AliSim-HPC is flexible: it can run on a personal computer with multi-threading, as well as on a distributed-memory cluster with many CPUs and multiple cores per CPU.

Our contributions are fourfold. First, this is the very first-time high-performance computing techniques are applied to phylogenetic sequence simulators. Second, we provide AliSim-HPC as an extension of IQ-TREE (Nguyen *et al*., 2015; Minh *et al*., 2020), an open-source and widely used phylogenetic software, thus, maximizing its usage and benefit to the user community. Third, we demonstrate that AliSim-HPC can efficiently simulate large genomic data sets, thus, facilitating large-scale benchmarking of phylogenetic methods and providing training data for machine learning-based applications. And fourth, we provide practical recommendations on the choice of the number of threads per process and multi-threading algorithms for simulating large genomic data sets.

## Materials and Methods

### The Sequential AliSim Algorithm

Here, we provide a brief summary of the published AliSim algorithm (Ly-Trong *et al*., 2022). Assuming that we want to simulate an alignment with *N* sequences, each of which contains *L* sites from a phylogenetic tree *T* and a substitution model *M*. Let *S*_!_ denote the sequence at node *j* of tree *T*. AliSim first generates a sequence with *L* sites at the root of the tree based on the state frequencies of model *M* (Fig. 2). Then, AliSim traverses tree *T* in a preorder manner to simulate a new sequence at each node based on the sequence of its parent node and the substitution model *M* (Ly-Trong *et al*., 2022). At tips, AliSim writes the simulated sequences to an MSA file. To generate many alignments, AliSim repeats this process sequentially. We present the sequential AliSim in the following.

**Fig. 2.**
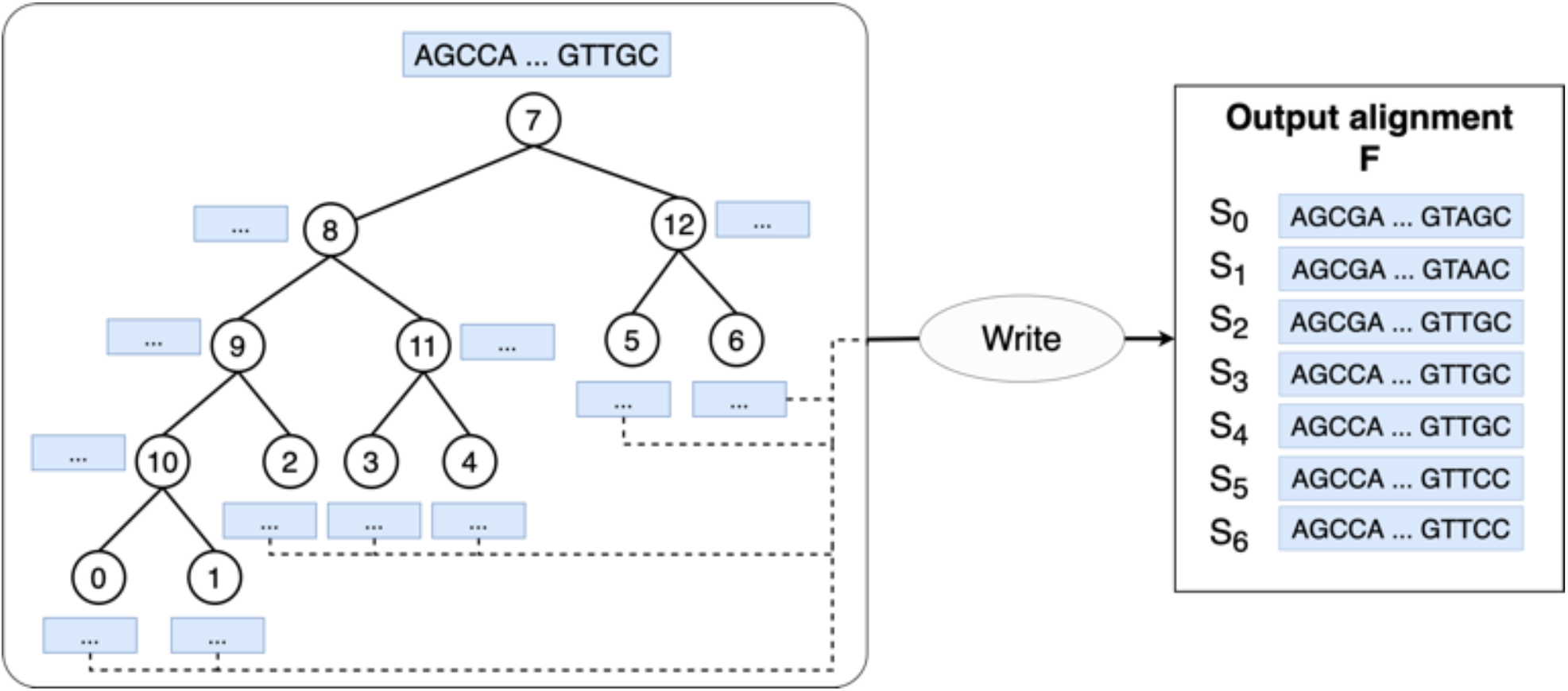
Illustration of the sequential AliSim algorithm to simulate a single alignment along an example tree. It starts by generating a sequence *S*_7_ at the root node 7, and subsequently, performs a preorder traversal visiting nodes 8, 9, 10, 0, 1, 2, 11, 3, 4, 12, 5, and 6. AliSim simulates *S*_8_ based on *S*_7_ and the substitution model *M*, and so on. At tip nodes 0 to 6, it writes *S*_0_ to *S*_6_ to an output alignment.

#### Algorithm 1: SequentialAliSim(*T, M, L, F*)

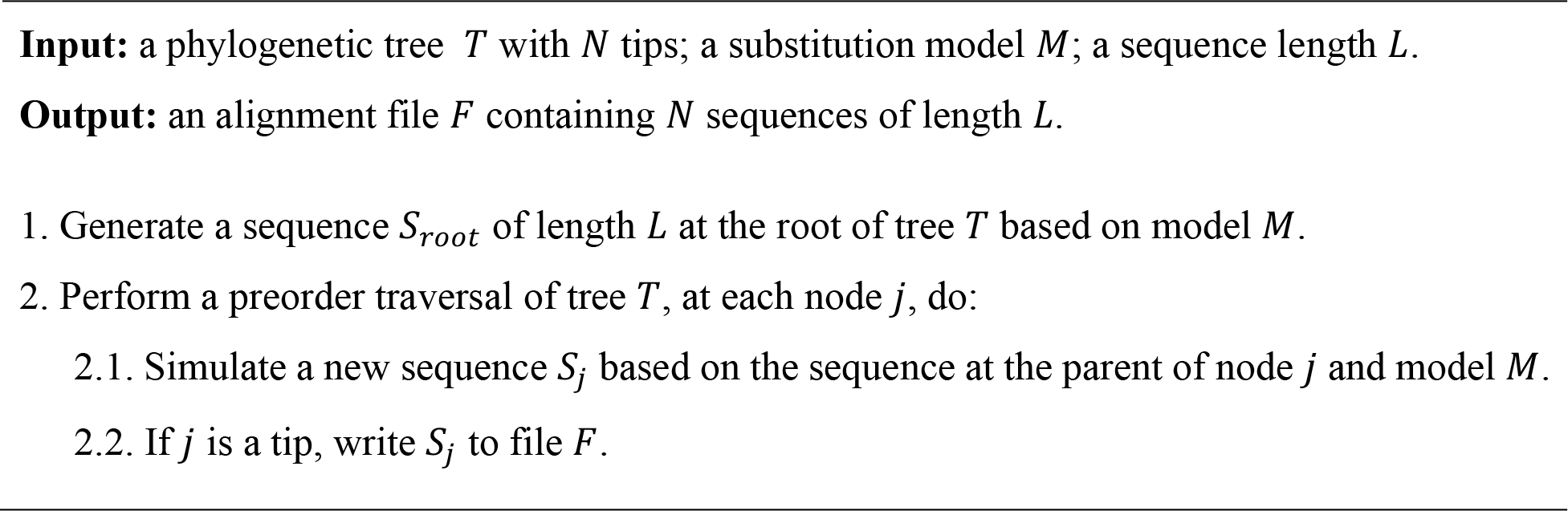

Thanks to a memory saving technique (Ly-Trong *et al*., 2022), the memory complexity of the sequential AliSim algorithm is *O(N) + O(D* ∗ *L)*, where *D* is the depth of tree *T*. The first *O(N)* and the second *O(D* ∗ *L)* terms represent the memory to store the tree structure and the simulated sequences, respectively.

### AliSim-OpenMP

We now introduce two different algorithms to parallelize AliSim using OpenMP. Most substitution models assume independence among sites. That naturally allows a straightforward parallel scheme for simulating an alignment with OpenMP (Fig. 3): each thread independently simulates a continuous block of the MSA with a roughly similar length. Given *k* threads, each thread *i* simply executes SequentialAliSim 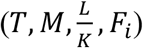 to generate a temporary file *F*_*i*_ containing *N* sequences of length 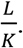.

**Fig. 3.**
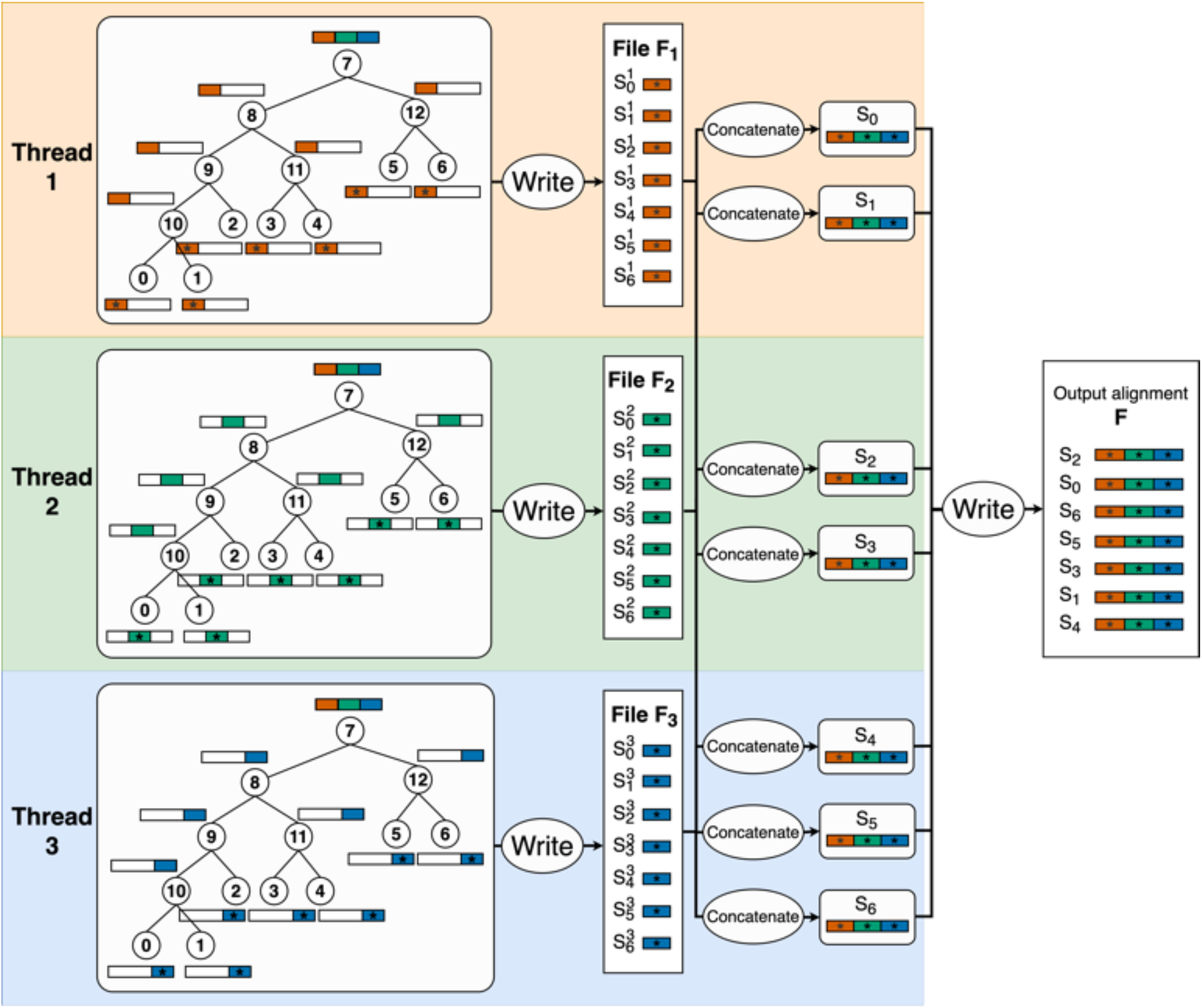
Illustration of the AliSim-OpenMP algorithm using external memory to simulate an alignment with 3 threads. These threads execute the sequential AliSim algorithm independently to generate 3 temporary files *F*_1-_, *F*_2_, *F*_3_; each file contains 7 subsequences of length ^)^. Then, thread 1 reconstructs two sequences *S*_0_ and *S*_1_ by concatenating their subsequences from all 3 temporary files. At the same time, thread 2 reconstructs *S*_2_ and *S*_3_ while thread 3 reconstructs *S*_4_, *S*_5_ and *S*_6_. The concatenated sequences are written one by one to the output alignment.

Next, we need to combine individual *F*_*i*_ files into a single alignment file *F*. AliSim invokes another parallel section, where each thread reads a subset of roughly 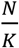 sequences across all temporary files, concatenates the subsequences into the full sequences of length *L*, then writes the full concatenated sequences to file *F* in a critical OpenMP section because file writing operations are not thread-safe. This algorithm is called “AliSim-OpenMP using external memory” because it creates temporary files to store intermediate alignments and is outlined below.

#### Algorithm 2: AliSimOpenMP_EM(*T, M, L, F, K*)

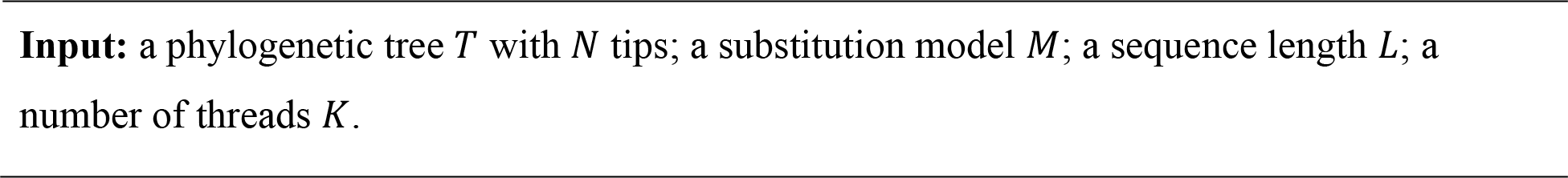

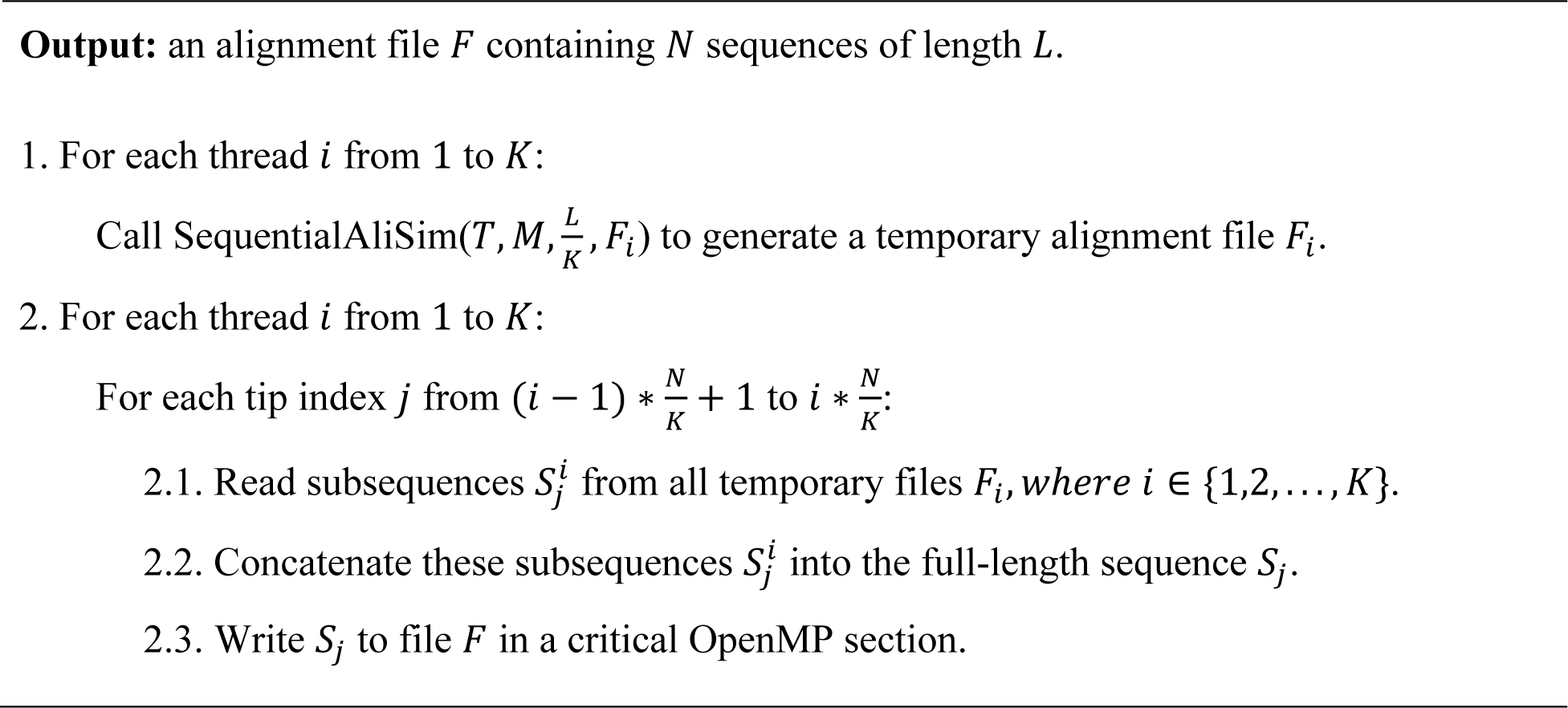

Algorithm 2 has the same memory complexity of *O(N) + O(D* ∗ *L)* as the sequential AliSim algorithm, where *D* is the depth of tree *T*. However, it consumes double the amount of external memory to store temporary files.

Algorithm 2 contains two parallel sections. The first section is embarrassingly parallel without any inter-thread communications, thus, we expect this section to gain linear speedup. However, the second section can be the main bottleneck due to too many I/O operations. A quick solution to deal with this problem would be to re-implement *F*_*i*_ as an internal memory storage. However, it requires an additional memory of *O(N* ∗ *L)*, which by large exceeds *O(D* ∗ *L)* and is therefore undesirable for large alignment simulations.

Therefore, we designed another algorithm called “AliSim-OpenMP using internal memory” (Fig. 4) to avoid writing temporary files as follows. We allocate *k* − 1 threads, each thread simulates one of the (*k* − 1) blocks of the MSA by calling a modified version of SequentialAliSim 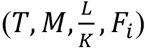 where *F*_*i*_ is now redesigned as a buffer in the internal memory. Whereas the last thread *k* is dedicated to only writing the buffers into the output file *F*.

**Fig. 4.**
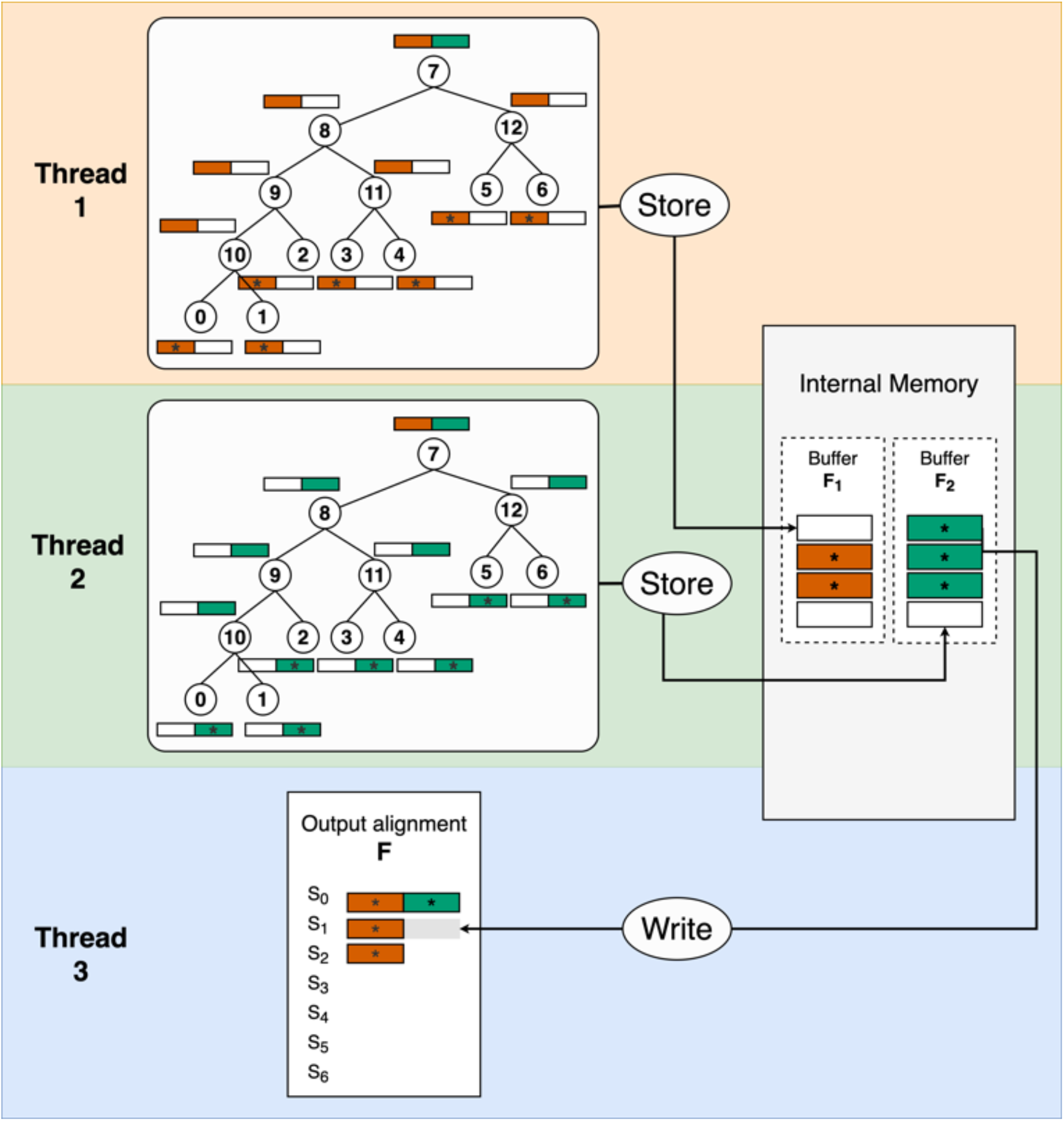
Illustration of the AliSim-OpenMP algorithm using internal memory to simulate an alignment with 3 threads. Threads 1 and 2 execute a modified version of the sequential AliSim algorithm independently to simulate subsequences of *S*_0_ to *S*_6_ of length 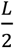 and store these subsequences in the corresponding buffers *F*_1_ and *F*_2_. Thread 3 continuously accesses buffers *F*_1_ and *F*_2_ to write subsequences into the right position of the output alignment *F* and then free the corresponding memory in the buffers. Thread 3 repeats that process until threads 1 and 2 are finished and all subsequences are outputted to the output alignment.

To reduce the RAM consumption, each thread does not store all *N* subsequences in its buffer *F*_*i*_, but only a fraction *N* ∗ *λ* of subsequences, where *λ* is a parameter between 0 and 1. Whenever each “simulating” thread *i* (from 1 to *k* − 1) simulated a subsequence 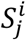 at tip *j*, it will store 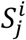 to *F*_*i*_ if *F*_*i*_ has some free memory. Otherwise, thread *i* will need to wait until *F*_*i*_ becomes available.

The I/O thread *k* continuously monitors the buffers. Whenever there is any subsequence 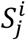 stored in any buffer *F*_*i*_, it will write 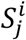 into the right position of file *F* and free the corresponding memory in *F*_*i*._ The I/O thread then checks the next buffer in the round-robin fashion (either *F*_*i+*1_ if *i* < *k* − 1 or *F*. if *i* = *k* − 1). This ensures a relative balance in memory availability among the buffers. This algorithm is outlined in Algorithm 3.

#### Algorithm 3: AliSimOpenMP_IM(*T, M, L, F, K*)

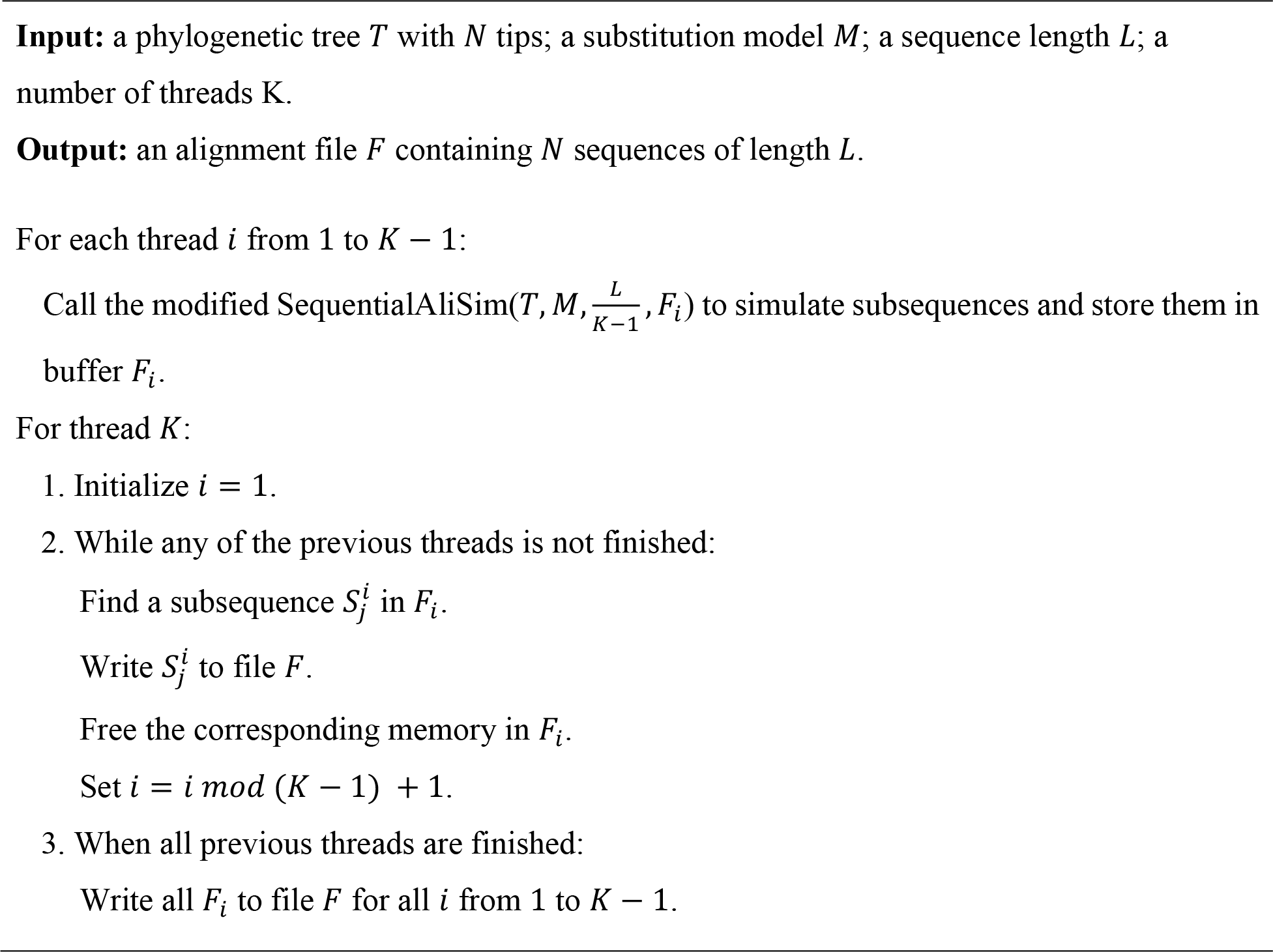

The memory complexity of this algorithm is *O(N) + O(D* ∗ *L + N* ∗ *λ* ∗ *L)*. Small *λ* will increase the waiting time of the “simulating” threads, thus potentially increasing the runtime. Whereas large *λ* will increase the RAM consumption. To balance the trade-off between runtime and RAM consumption, we set the default *λ* to *min((k* − 1) ∗ 2/*N*, 1) because with more threads (higher *k*), each thread needs to simulate shorter subsequences, which is faster than having fewer threads, and therefore a larger buffer size is needed.

The two AliSim-OpenMP algorithms introduced above have their own advantages and disadvantages depending on the simulating conditions, but they will complement each other.

### AliSim-MPI

A more practical demand is to simulate many alignments. Therefore, we developed AliSim-MPI to embarrassingly parallelize those simulations across a distributed-memory system. Specifically, to simulate *H* alignments using *P* MPI processes, AliSim-MPI simulates roughly 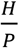 alignments per process (Fig. 5). These processes perform simulations independently and write separate alignment files. No communication is needed between the processes. The memory complexity is proportional to the number of processes: *O(P* ∗ *N) + O(P* ∗ *D* ∗ *L)*.

**Fig. 5.**
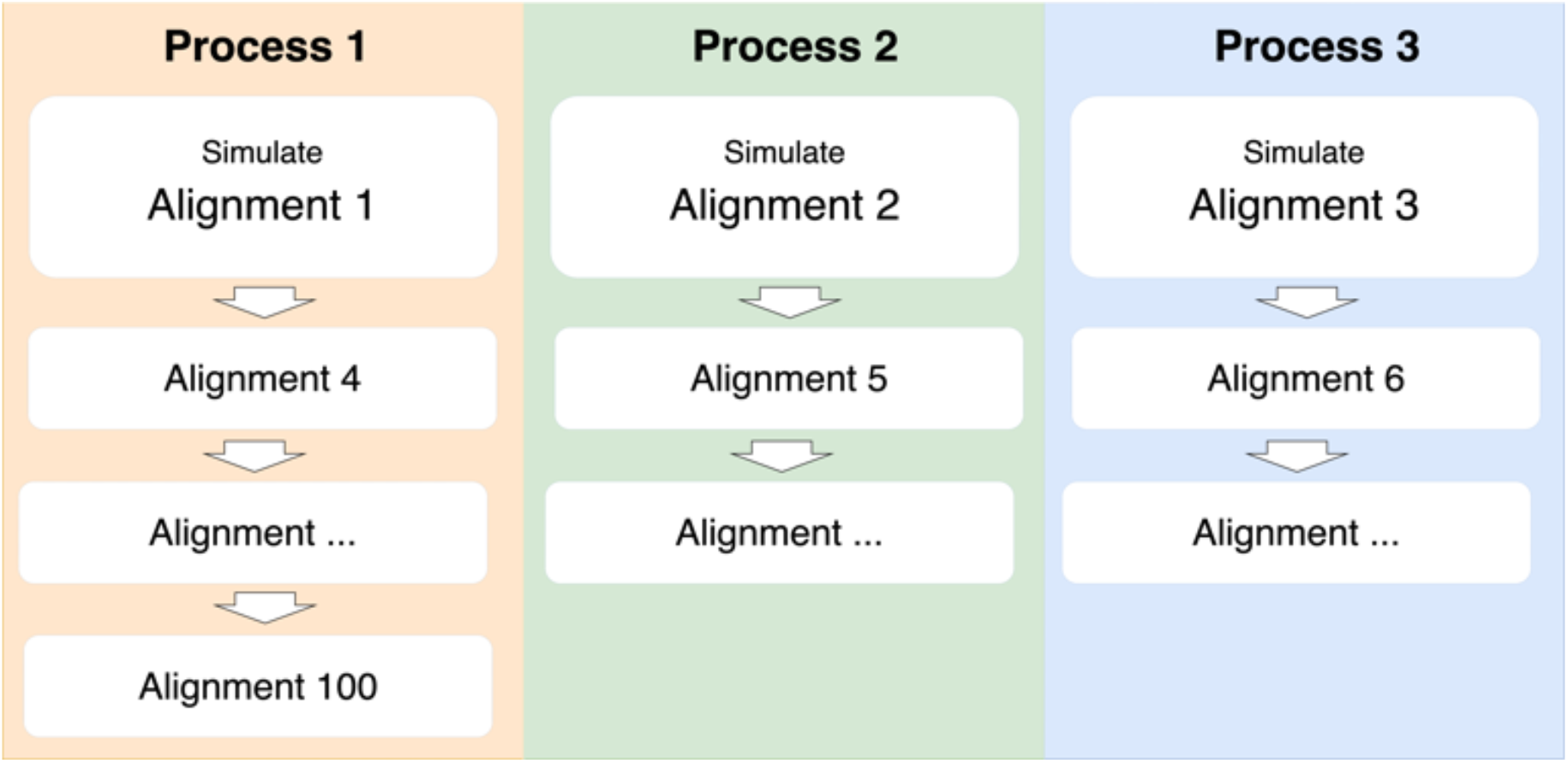
Illustration of the AliSim-MPI algorithm to simulate 100 alignments with 3 processes. Process 1 simulates 34 alignments 1, 4, …, and 100. Whereas processes 2 and 3 simulate 33 alignments per process. These processes run independently without any inter-process communications.

### AliSim-HPC for high-performance computing systems

We now combine AliSim-OpenMP and AliSim-MPI to enable simulations on a large cluster with *P* processes, each having *k* threads (i.e., a total of *P* ∗ *k* threads are run in parallel). The AliSim-HPC algorithm is outlined below.

#### Algorithm 4: AliSimHPC(*T, M, L, H, P, K*)

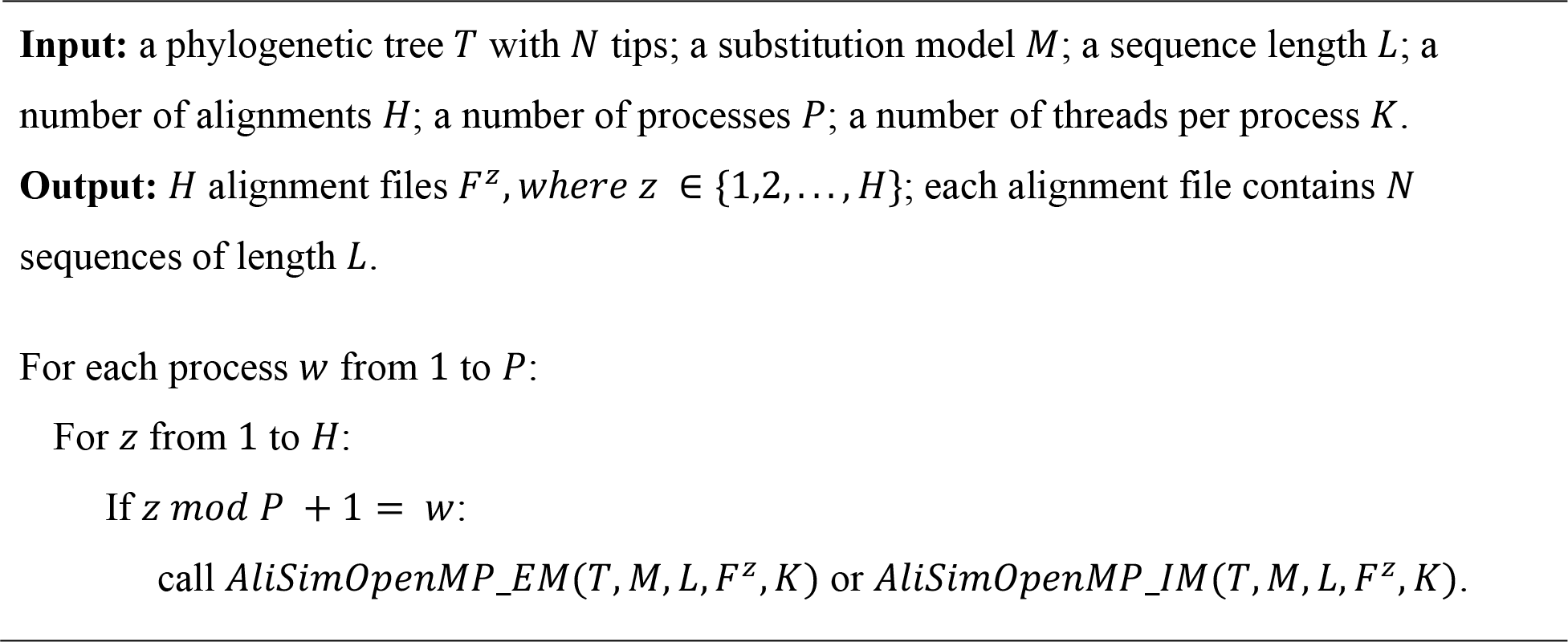

We thereby refer to the two variants of AliSim-HPC that integrate AliSim-OpenMP using external memory and internal memory as AliSim-HPC-EM and AliSim-HPC-IM, respectively.

The memory complexity is *O(P* ∗ *N) + O(P* ∗ *D* ∗ *L)* for AliSim-HPC-EM; and *O(P* ∗ *N) + O(P* ∗ *D* ∗ *L + P* ∗ *N* ∗ *λ* ∗ *L)* for AliSim-HPC-IM.

### Random generator initialization

Reproducibility is strongly desirable in any software, which involves random number generation. To ensure this, AliSim-HPC employs The Scalable Parallel Random Number Generators Library (SPRNG) (Mascagni and Srinivasan, 2000) and requires users to specify a random number generator seed *r*. It then computes a unique seed number for each thread of each process as *r + p* ∗ 1000 *+ k*, where *p* and *k* denote the process and thread IDs, respectively. If *r* is not provided, it will be set to the current microsecond of the CPU.

### Benchmark setup

We evaluated the performance of the two variants of AliSim-HPC (using external and internal memory) compared with the sequential AliSim on the Gadi supercomputer (NCI’s newest supercomputer is Gadi, Australia’s peak research supercomputer for 2020 and beyond), a cluster of 3,200 nodes with a total of 155,000 CPU cores, 567 TB of RAM, and 640 GPUs. We employed up to six computing nodes, each of which has 2 × 24-core Intel Xeon Platinum 8274 (Cascade Lake) 3.2 GHz CPUs and 400 GB SSD. We run all experiments with Open-MPI v4.1.3 (Open MPI: Open Source High Performance Computing) and OpenMP 4.5.

We measured the strong scaling behavior and total RAM consumption when simulating 100 large alignments. Due to storage quota and resource capacity per node, we set the maximum number of processes and the number of threads per process at 32. By varying the number of processes *P* and the number of threads per process *k* at 1, 2, 4, 8, 16, and 32, we formed a total of 33 combinations, such that the total number of CPU cores (the number of processes times the number of threads) was up to 256. We simulated two types of large alignments: 1M sequences of 30K sites, which we called a ‘deep alignment’; and 30K sequences of 1M sites, which we called a ‘long alignment’. The input phylogenetic trees were randomly drawn under the Yule-Harding model (Yule, 1925; Harding, 1971) and exponentially distributed branch lengths with a mean of 0.1. For the model of evolutions, we applied the General Time Reversible (GTR) (Tavaré, 1986) with an invariant site proportion of 0.2 and discrete Gamma rate heterogeneity (Gu *et al*., 1995) with a Gamma shape of 0.5.

## Results

### Two AliSim-OpenMP algorithms complemented each other

We first benchmark the pure multi-threading algorithms (AliSim-OpenMP) without multi-processing. Fig. 6 shows the performance of the two AliSim-OpenMP algorithms using internal (IM) and external memory (EM). For long alignment simulations, when the number of threads is 8 or more, AliSim-OpenMP-IM obtained 7.6 – 10.2 fold speedups (compared with the sequential AliSim), while the speedups for AliSim-OpenMP-EM were slightly lower at 4.8 – 9.7 folds (Fig. 6A). In contrast, for deep alignment simulations, AliSim-OpenMP-EM obtained higher speedups (from 1.7 to 8.8 folds) than the IM variant (from 1.1 to 3.2 folds) (Fig. 6C). Therefore, the two versions complemented each other.

**Fig. 6.**
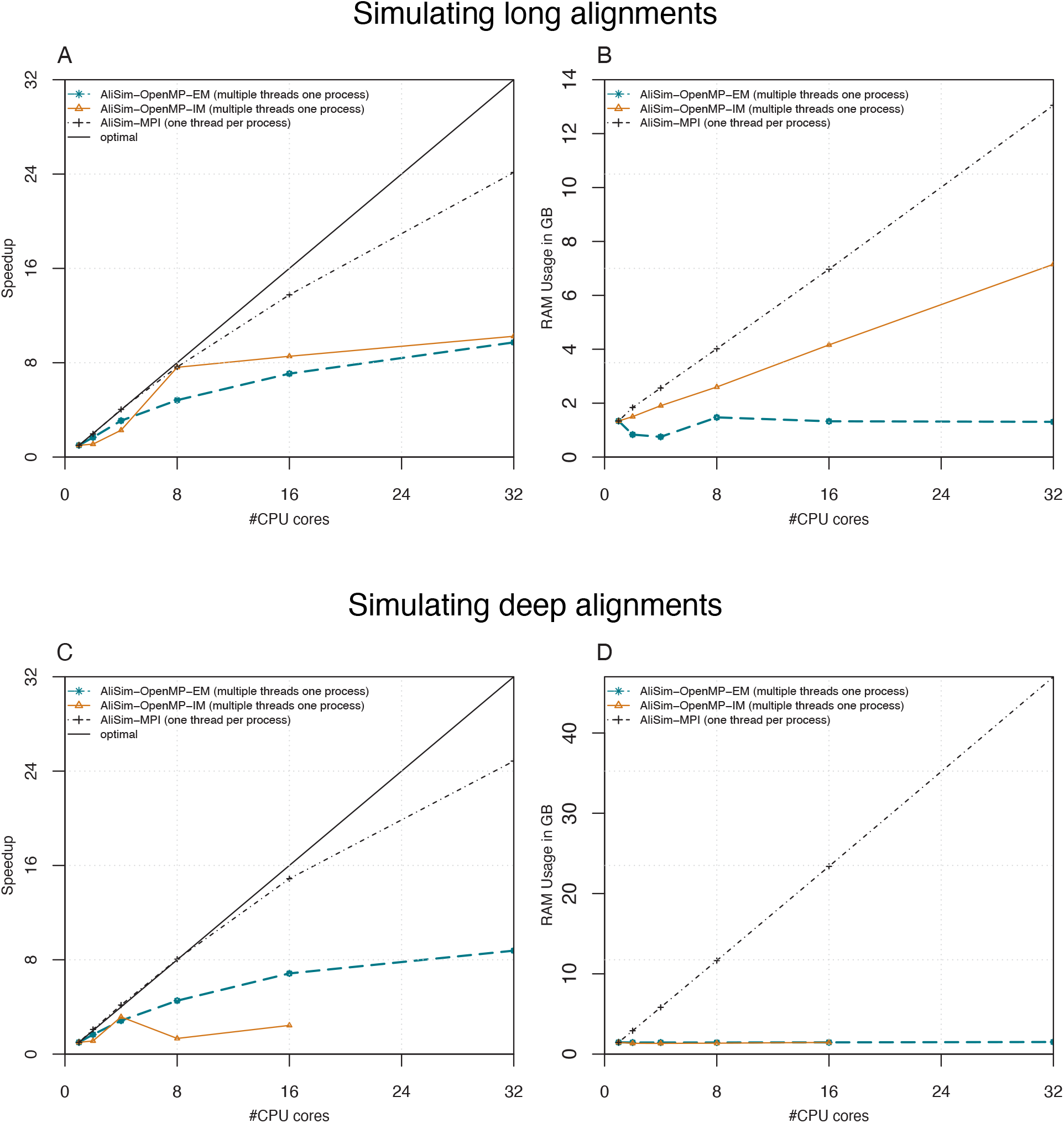
Strong scalings and peak RAM consumptions of the two AliSim-OpenMP algorithms, and AliSim-MPI for long-alignment (30K sequences x 1M sites; sub-panels A and B); and deep-alignment (1M sequences x 30K sites; sub-panels C and D) simulations. In deep-alignment simulations, AliSim-OpenMP-IM using 32 threads per process took an excessively long runtime, thus we skipped that test to save the computational resources.

Regarding the RAM consumption, AliSim-OpenMP-IM, and AliSim-OpenMP-EM, as expected, consumed 1.3 – 7.2, and 1.3 – 1.5 GB RAM, respectively, to simulate long alignments (Fig. 6B). Interestingly, the peak RAM consumptions of the two AliSim-OpenMP algorithms were almost identical at around 1.5 GB RAM regardless of the number of threads (Fig. 6D) for deep simulations. That is because the memory complexity of both versions (see Materials and Methods) is proportional to the alignment length *L*, which is more noticeable in the long than the deep simulations.

### AliSim-MPI obtained high parallel efficiency

Next, we benchmark the pure multi-processing AliSim-MPI version, where each process is single-threaded. AliSim-MPI achieved almost linear speedup with high parallel efficiency (Fig. 6A and C). With 32 CPU processes, AliSim-MPI achieved approximately 25-fold speedup compared with the sequential AliSim, a 78% parallel efficiency for both long and deep alignment simulations.

The RAM consumption grew, as expected, proportionally with the increasing number of processes (Fig. 6B and D). AliSim-MPI required 1.3 – 13 GB, and 1.5 – 47 GB RAM to simulate long and deep alignments, respectively. Due to a large number of nodes *N* in the phylogenetic tree, simulating deep alignments consumed more RAM than long alignments (see Materials and Methods).

### AliSim-HPC achieved excellent strong scaling behavior

We now benchmark AliSim-HPC that combines the benefits of AliSim-OpenMP (low RAM consumptions) and AliSim-MPI (excellent speedups). Fig. 7 illustrates the performance of the two variants AliSim-HPC-EM and AliSim-HPC-IM using external and internal memory, respectively. For long alignment simulations, while both variants achieved excellent strong scaling when increasing the total number of CPU cores (*P* ∗ *k*), AliSim-HPC-IM (Fig. 7B) obtained higher speedups than AliSim-HPC-EM (Fig. 7A). For example, AliSim-HPC-IM reached 162-fold speedup using 32 processes x 8 threads (Fig. 7B), but AliSim-HPC-EM only reached 90-fold speedup. In fact, AliSim-HPC-EM achieved 182-fold speedup in step 1 (of Algorithm 2) compared with the sequential AliSim algorithm. However, AliSim-HPC-EM required an additional phase (step 2 in Algorithm 2) to concatenate temporary files, which took approximately the same amount of the runtime of step 1, thus reducing the overall performance of that algorithm.

**Fig. 7.**
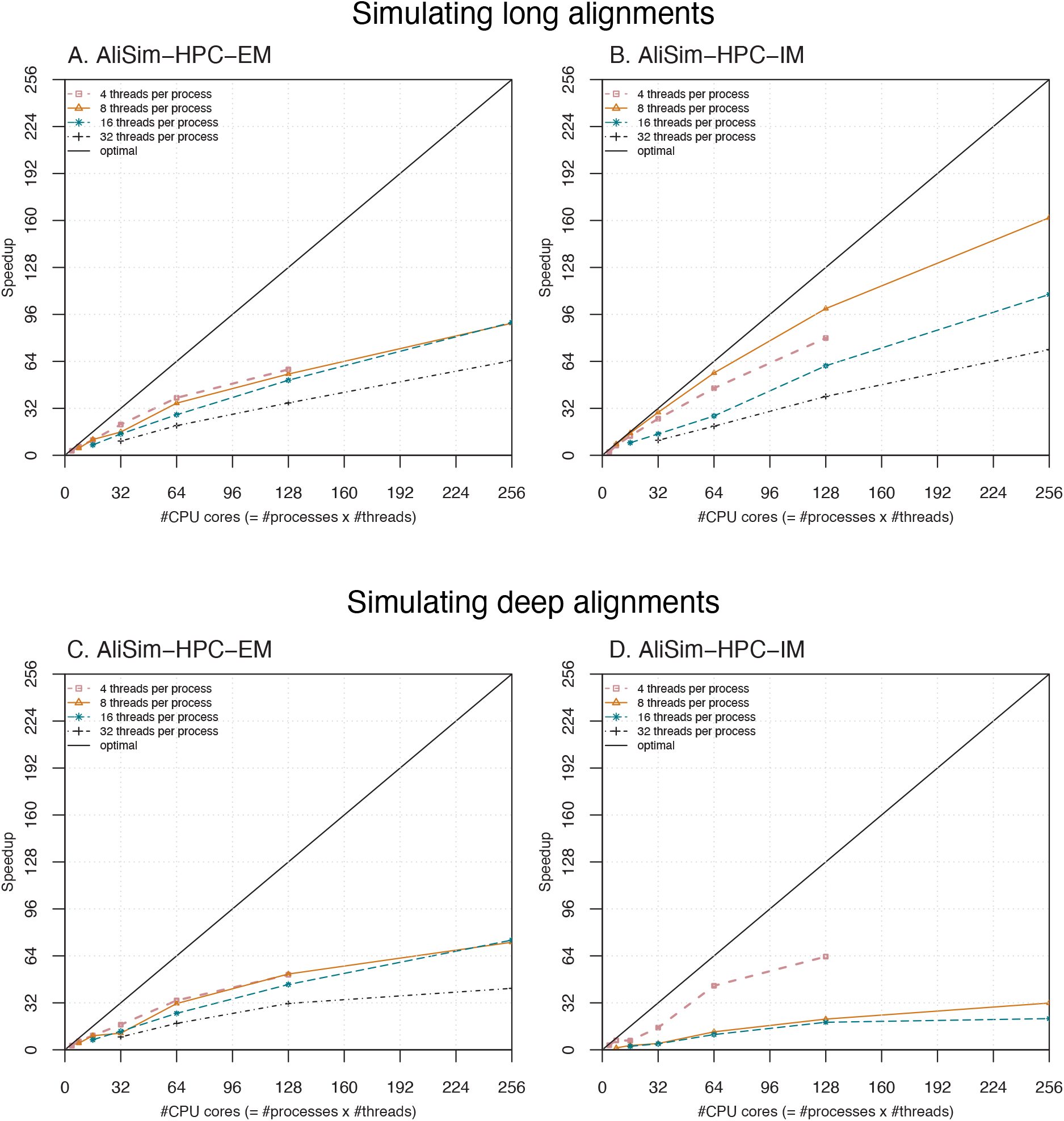
Strong scaling of AliSim-HPC-EM and AliSim-HPC-IM in long-alignment (sub-panels A and B); and deep-alignment (sub-panels C and D) simulations. In deep-alignment simulations, the curve of AliSim-HPC-IM using 32 threads per process is missing since these tests took excessively long runtime, thus we skipped them to save the computational resources.

For simulating deep alignments, AliSim-HPC-EM (Fig. 7C) often outperformed AliSim-HPC-IM (Fig. 7D). AliSim-HPC-EM obtained a 73-fold speedup compared with only 32-fold speedup of AliSim-HPC-IM for 32 processes x 8 threads. But interestingly, AliSim-HPC-IM with 4 threads per process performed better than the EM variant, obtaining 64-fold speedup for 32 processes x 4 threads compared with 51-fold speedup for AliSim-HPC-EM. Unfortunately for this setting, we could not run our tests with a higher number of processes due to excessive memory requirements.

In summary, both versions of AliSim-HPC achieved excellent scaling behavior. The best setting of AliSim-HPC-EM and AliSim-HPC-IM reduced the wall-clock time from one day to 9 and 19 minutes for simulating long and deep alignments, respectively.

### The RAM consumption of AliSim-HPC increased with the number of processes

Fig. 8 shows the memory footprint of AliSim-HPC-EM and AliSim-HPC-IM. As expected, the RAM consumption of the two variants increased with the number of processes *P*.

**Fig. 8.**
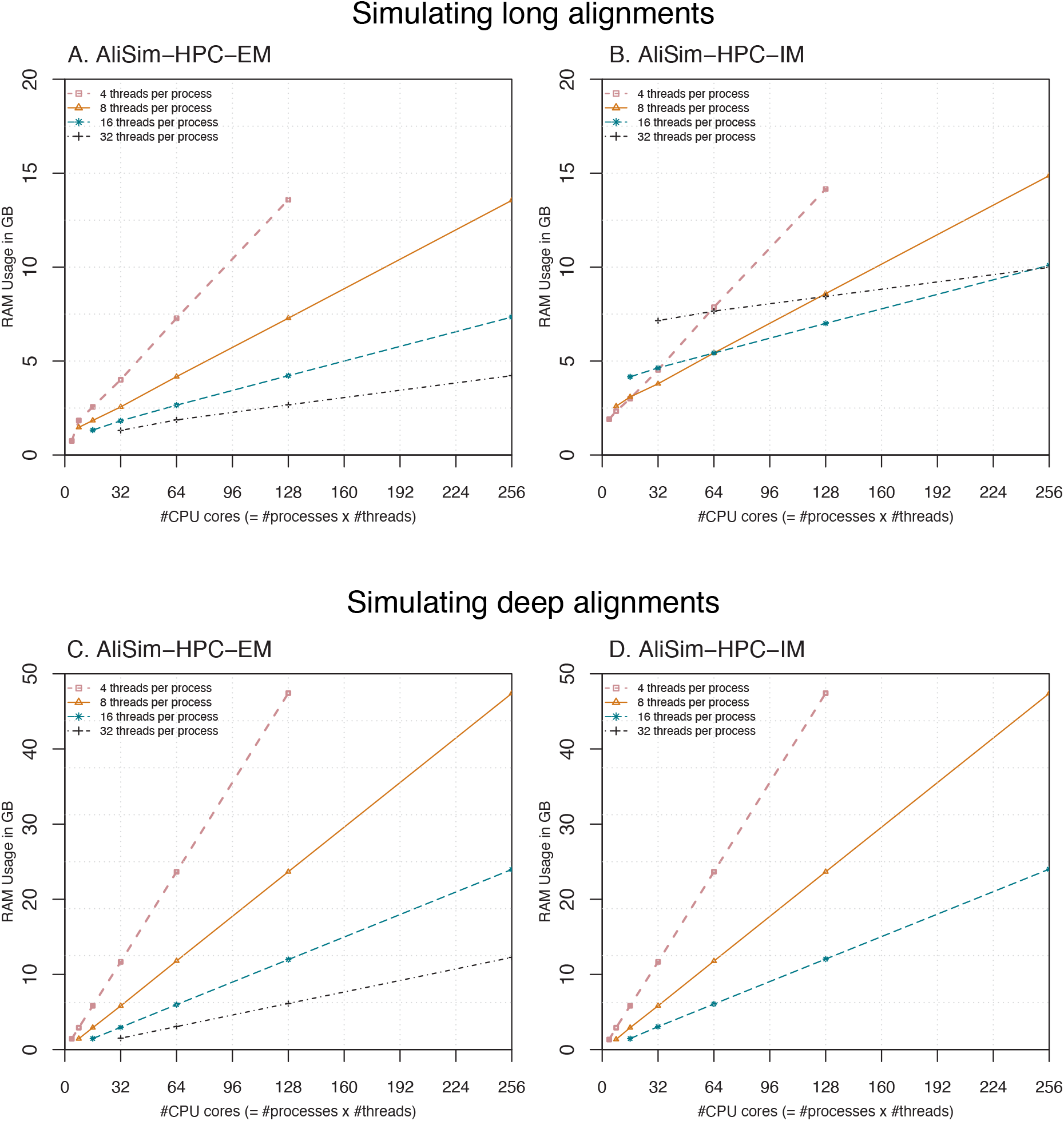
The memory footprint of AliSim-HPC-EM and AliSim-HPC-IM in long-alignment (sub-panels A and B); and deep-alignment (sub-panels C and D) simulations.

For long-alignment simulations, with 8 threads per process, AliSim-HPC-EM consumed 1.5 -

13.6 GB RAM when increasing the number of processes from 1 to 32 (Fig. 8A). Whereas, AliSim-HPC-IM took 2.6 -15 GB RAM for the corresponding settings (Fig. 8B).

For deep-alignment simulations, the memory footprints of the two variants of AliSim-HPC were almost identical (Fig. 8C and D). Using 32 processes, they consumed approximately 47 GB RAM regardless of the number of threads per process. With the same number of processes, the two variants of AliSim-HPC consumed the same amount of RAM as AliSim-MPI.

## Conclusion and Future Work

This paper introduces AliSim-HPC, a high-performance sequence simulator for phylogenetics. We present two multi-threading algorithms to simulate a single large alignment with OpenMP, and an embarrassingly parallel scheme to simulate many alignments with MPI on a distributed-memory system. In the future, we would also like to extend AliSim-HPC to employ SIMD (Cardoso *et al*., 2017) and GPU-based parallelization.

With an appropriate setting of the number of threads per process, AliSim-HPC is highly efficient because it involves minimal inter-thread and no inter-process communications. Strong scaling experiments also showed that AliSim-HPC is scalable: we obtained 162-fold speedup when employing 256 CPU cores to simulate 100 large alignments; further speedup is achievable with more computational resources since the parallelization is not yet saturated.

Among all settings, the 8 threads per process likely yielded the best performance. However, we observed that if using AliSim-HPC-EM, two settings, 8 threads and 16 threads per process, performed equally well. Therefore, we recommend applying 16 threads per process since it can save half of the memory consumption compared with the 8 threads per process setting.

The performance of the two AliSim-HPC variants greatly depends on the sequence length. Based on these experimental results, we recommend applying AliSim-HPC-IM for simulating long alignments. In contrast, to simulate short and moderate sequences (e.g., 30K sites), AliSim-HPC-EM is preferable.

However, determining whether an alignment is long or short is subjective and the performance of our algorithms also relies on the hardware (e.g., processor, SSD/HDD storage). Therefore, our future work covers designing a mechanism to automatically select the best algorithm on the fly.

The design of AliSim-OpenMP is based on the assumption that different sites in the alignment evolve independently. However, this assumption does not hold for some advanced models, such as insertion-deletion. In the case of site non-independent models, a different parallel strategy is needed. A feasible idea is to perform level order traversal on the tree so that sequences at same-depth nodes can be simulated simultaneously; each thread simulates a full-length sequence at a node. Besides, if the number of alignments is much greater than the number of CPU cores, we can also extend our AliSim-OpenMP algorithms so that each thread can simulate entire alignments independently to avoid the writing bottleneck.

Finally, the I/O operations are currently the bottleneck of our AliSim-OpenMP algorithms, which explains their far-from-perfect speedups (Fig. 6). Future work would employ parallel I/O techniques that will remove this bottleneck and make AliSim-HPC even more efficient for much larger scale simulations.

## Data Availability

The data underlying this article are available in the Zenodo Repository at https://doi.org/10.5281/zenodo.7407513.

## Acknowledgements

This research was undertaken with the assistance of resources and services from the National Computational Infrastructure (NCI), which is supported by the Australian Government. We thank Robert Lanfear, Thomas Wong, James Barbetti, Fred Jaya, Huaiyan Ren, Jeremias Ivan, Caitlin Cherryh, and Matthew Macaulay for their valuable comments and discussions.

